# CARS: A General Force Field for Carotenoids

**DOI:** 10.64898/2026.07.31.742033

**Authors:** Andrey Nikolaev, Yaroslav Orlov, Victoria Khanina, Ivan Gushchin

## Abstract

Carotenoids are structurally diverse isoprenoid pigments that play central roles in photosynthesis, photoprotection, membrane organization, and cellular signaling. Despite their biological and technological importance, atomistic simulations of carotenoids remain limited by the lack of a transferable force field spanning the chemical diversity of naturally occurring compounds, including glycosylated and acylated derivatives. Here we present CARS (CARotenoidS), a transferable force field for carotenoids that integrates seamlessly with the AMBER family of biomolecular force fields. Parameters were systematically optimized against 22 957 r2SCAN-3c reference energies for 25 representative molecular fragments, yielding an accurate description of polyene conformational energetics, ring rotations, and molecular geometries. Across diverse validation sets, CARS substantially outperforms GAFF2 and provides improved agreement with quantum-chemical reference data for glycosylated and acylated carotenoids. For zeaxanthin, CARS also surpasses OPLS-AA, CGenFF, and previously published carotenoid-specific parameters set in reproducing conformational energetics and structural properties. Two complementary parameter sets are provided: CARS for glycosylated and non-lipidated carotenoids, and CARS+Lipid21 for carotenoids containing saturated or monounsaturated lipid chains. By providing the first unified and transferable parameterization covering the structural diversity of natural carotenoids while remaining fully compatible with established AMBER force fields, CARS removes the need for molecule-specific reparameterization and enables reliable molecular simulations of carotenoids in proteins, membranes, and other complex biological assemblies.

## Introduction

Carotenoids are isoprenoid pigments synthesized by plants, algae, cyanobacteria, and some fungi, and they are acquired by animals through diet. In photosynthetic organisms, they are involved in light harvesting and photoprotection; in non-photosynthetic tissues, they function as antioxidants and as precursors of signalling molecules^1^. This broad biological distribution is accompanied by substantial chemical diversity (more than 1100 species are described^2^), which allows carotenoids to operate in membranes, pigment–protein complexes, and aqueous environments.

All carotenoids share a conjugated polyene chain that determines their protochemical properties and their overall geometry^3^. The chain is capped by end groups, most often beta-rings, epsilon-rings, or acyclic psi-ends (Figure 1). This core structure is frequently modified by hydroxylation, epoxidation, ketolation, and introduction of allenic or acetylenic bonds. A further layer of diversity arises from covalent attachment of sugars or fatty acids, which results in glycosylated and acylated derivatives. These modifications alter the polarity, amphiphilicity, and aggregation behaviour of carotenoids, and determine how carotenoids are positioned in lipid bilayers and recognised by proteins.

**Figure 1.**
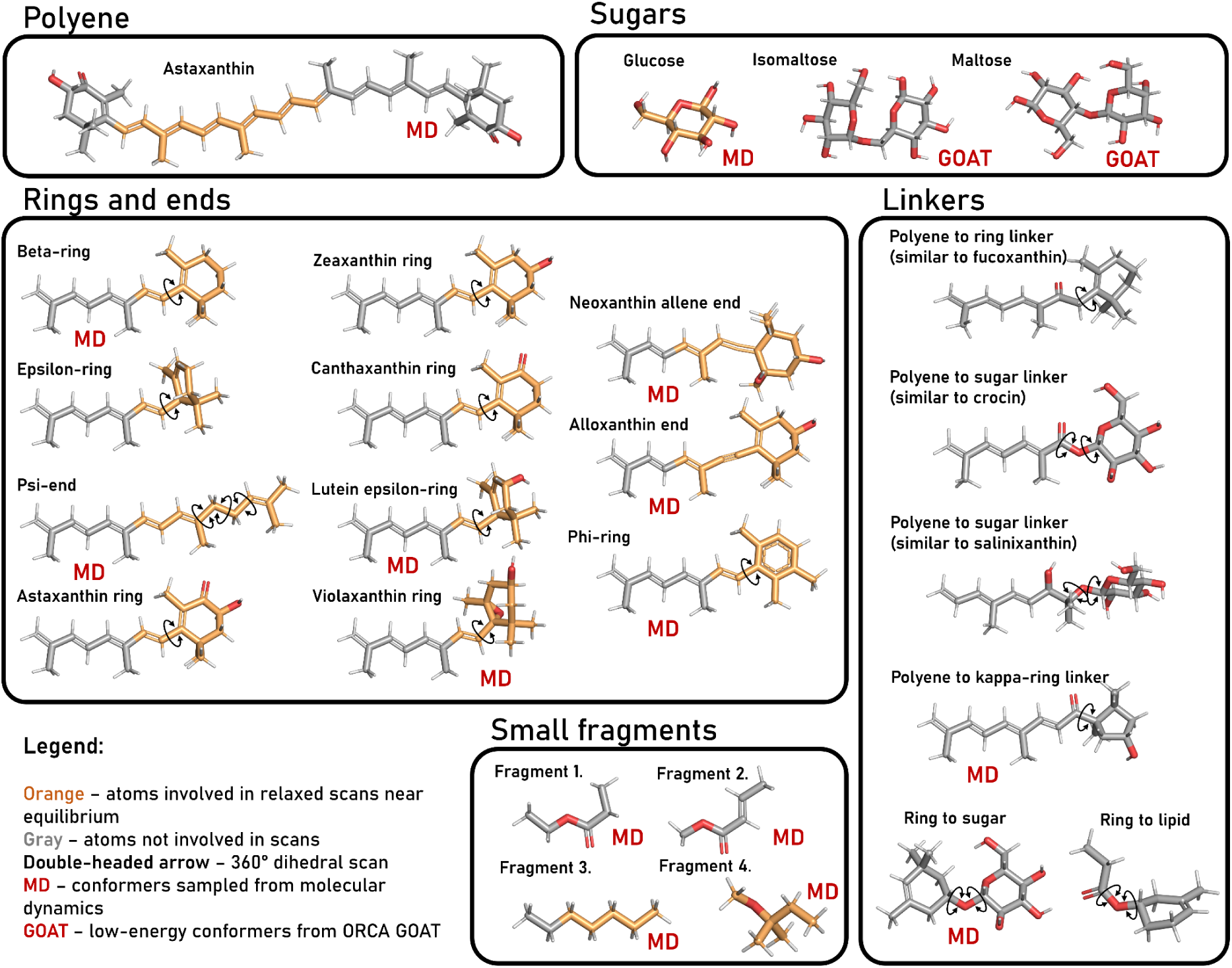
Training set for the CARS force field parameterization. Each molecule was perturbed using one or more deformation strategies. Atoms involved in relaxed surface scans around the equilibrium values of internal coordinates (bond lengths, angles, and dihedrals) are highlighted in orange. Double-headed arrows indicate dihedral angles scanned over a full 360° rotation. Molecules with the MD label were additionally sampled through molecular dynamics simulations in water at 300 K. Molecules with the GOAT label were subjected to low-energy conformational sampling using the GOAT procedure as implemented in ORCA^35,58^.

Reliable MD simulations of carotenoid-containing systems require parameters that accurately describe the conjugated polyene, end-group rotations, and covalent modifications. Lipid-specific parameter sets such as Lipid21^4^ and CHARMM36^5^ are trained on saturated and monounsaturated chains; therefore, their training sets do not include the extended conjugation of carotenoid polyenes. General-purpose force fields require validation before application to carotenoids. For GAFF^6^, tests on pyridine revealed substantially underestimated barriers for polyene isomerisation^7^. OPLS-AA^8^ has been applied to study several xanthophylls including lutein and zeaxanthin and yielded acceptable agreement with experimental data^9^, but its performance across the wider chemical space of carotenoids is not established.

These gaps motivated development of specialized parameters for individual carotenoids. Early work produced a CHARMM36-compatible set for beta-carotene and zeaxanthin using B3LYP/6-31G(d)^10,11^ and MP2/6-31G(d)^12^ as the quantum mechanics (QM) reference^13^ and AMBER-compatible parameters for beta-carotene using B3LYP/6-31G* as the QM reference^14^. A broader contribution came from the work by Prandi et al., in which parameters for four xanthophylls (neoxanthin, violaxanthin, lutein, and zeaxanthin) were developed employing B3LYP/6-31G(d,p) as the reference^15^. This parameter set served as the starting point for parametrisation of other carotenoids: echinenone^16^, canthaxanthin^17^, okenone^18^, and salinixanthin^19^, for which parameters were developed by combining the parameters developed by Prandi et al. with GAFF parameters and refitting selected torsional terms. Recently, we developed parameters for astaxanthin based on GAFF2 and custom parameters for torsion terms in polyene, which were fitted using B3LYP-D3/def2-SVP^20,21^ as a reference^22^.

Parameterisation of carotenoids relies on QM reference data. However, the large size of carotenoids makes high-level calculations such as CCSD(T)^23^ with the complete basis set (CBS)^24^ extrapolation prohibitively expensive, forcing developers to rely on modest levels of theory such as B3LYP with small basis sets. This constraint affects the quality of the training data and, ultimately, the accuracy of the developed force fields. The emergence of a new wave of composite QM methods which combine high accuracy and low computational cost promises a solution to this problem^25–28^.

In this work, we describe the construction of a large training set containing deformation energies for 25 molecular fragments that cover the polyene chain, the most common carotenoid end-groups, as well as sugars, lipids, and the linkages between them. Using this training set, we develop CARS (CARotenoidS), a general force field for carotenoids that combines parameters from GAFF2 and Lipid21 with parameters trained in this work. We show that CARS accurately reproduces polyene torsional stiffness, ring-rotation energy profiles, and carotenoid geometries. It substantially surpasses GAFF2 in predicting deformation energies of glycosylated and acylated carotenoids. Because CARS is built in the Amber format, it can be used as easily as GAFF2, and alongside GAFF2, in standard simulation workflows. The CARS force field thus makes accurate simulations of chemically diverse natural carotenoids broadly accessible.

## Results

### Reference QM methods

We selected the r2SCAN-3c method for all single-point energy evaluations^28^. On the large GMTKN55 database, r2SCAN-3c achieves a WTMAD-2^29^ (weighted total mean absolute deviation relative to CCSD(T)/CBS) of 7.5 kcal/mol, close to the popular hybrid functionals B3LYP-D4 (6.5 kcal/mol) and PBE0-D4 (6.6 kcal/mol) with the accurate D4 dispersion correction^30^ and large aug-def2-QZVP basis set, as well as MP2/def2-QZVPP (6.9 kcal/mol^31^), but at a computational cost that is 100–1000 times lower. This speedup is essential because the training set requires tens of thousands of single-point calculations on diverse molecules.

We also used r2SCAN-3c to optimize the reference geometries of selected carotenoids, providing a benchmark for assessing geometry optimization quality across different force fields. However, building the training set required several thousand relaxed scans of internal coordinates for various molecules. For this task, we employed GFN2-xTB^32^, which reliably predicts geometries of organic molecules. The energies of the relaxed structures were subsequently recalculated at the r2SCAN-3c level, because GFN2-xTB energies are not sufficiently accurate (WTMAD-2 of approximately 25 kcal/mol^33^).

### Construction of the training set

Construction of the training set began with selection of small molecules representing fragments of naturally occurring carotenoids (Figure 1). The set includes the following end groups: beta-ring, epsilon-ring, phi-ring, and an open-chain psi-end. To preserve the chemical context, each ring was modeled together with a conjugated chain segment corresponding to one half of the full polyene backbone. The set also contains rings of the most common xanthophylls: canthaxanthin, zeaxanthin, astaxanthin, lutein, violaxanthin, neoxanthin, and alloxanthin. To enable the model to predict the properties of a complete conjugated polyene system, a full molecule of astaxanthin was included.

Glycosylation was represented by six-membered sugar molecules: glucose, maltose, and isomaltose. Lipid fragments were modeled using four small molecules: two fragments containing an ester bond (one of which includes a conjugated alkene double bond), a heptane molecule, and a branched lipid with an ether bond.

To capture interactions between different molecular fragments, the training set was supplemented with molecules representing several linkage types. These include: (1) a non-standard polyene–beta-ring linkage via a carbonyl group, analogous to fucoxanthin; (2) a polyene–sugar ester bond, analogous to crocin; (3) a polyene–sugar ether bond, analogous to salinixanthin; (4) a polyene–κ-ring linkage via a carbonyl group; (5) an ether bond between a beta-ring and a sugar; (6) an ester bond between a lipid and a sugar.

To generate a set of relevant molecular deformations for model training, three different approaches were employed.

In the first approach, a relevant fragment of the molecule was selected, and a relaxed surface scan of all internal degrees of freedom (bond lengths, valence angles, and dihedral angles) was performed around the equilibrium geometry. In a relaxed scan, the chosen coordinate is fixed at a series of values, while all other degrees of freedom are allowed to relax to their energy minimum at each step, yielding a physically meaningful deformation energy profile. Bond lengths were scanned within ±0.075 Å of the equilibrium value, valence angles within ±10°, and dihedral angles within ±30°. Each scan contained six points. Only the deformations with energy below 3 kcal/mol relative to the unstrained molecule were retained. For selected rotatable dihedral angles critical to the accurate description of carotenoid conformational dynamics, full 360° scans were performed.

In the second approach, conformations of selected molecules were sampled using molecular dynamics simulations. Simulations were run with the GAFF2 force field in explicit TIP3P^34^ water for 1 ns, and 1000 frames were extracted. For carotenoids, the GAFF2 force field was modified to include more physically accurate dihedral force constants for the polyene chain (see Methods), preventing non-physical isomerization.

In the third approach, the GOAT conformer search algorithm^35^, as implemented in the ORCA software package, was applied to sugars. Sugar geometries feature numerous local energy minima, which are difficult to sample adequately either by short molecular dynamics simulations or by systematic 360° dihedral scans. The GOAT algorithm performs a stochastic search to locate these minima efficiently.

Altogether, the three approaches yielded 22 957 reference QM energies (single-point r2SCAN-3c calculations) across all molecules and deformation types.

### Loss function

For loss evaluation, the energy of every structure in the training set was computed with the force field being trained. A separate loss term was calculated for each molecule and each deformation type. The total loss was defined as a weighted sum of these terms. Conformations sampled from molecular dynamics were assigned an 1 − R^2^ loss with a weight of 3. Relaxed scans of all internal coordinates around equilibrium were assigned an 1 − R^2^ loss with a weight of 0.7. Relaxed scans of dihedral angles around equilibrium were assigned the same 1 − R^2^ loss with a lower weight of 0.3. Full 360° relaxed scans of dihedral angles were assigned an MSE loss scaled by 1/(kcal/mol)^2^ with a weight of 0.1. Conformations sampled with the GOAT algorithm were assigned an 1 − R^2^ loss with a weight of 0.6. For astaxanthin, all contributions were multiplied by an additional factor of 3 to emphasize the importance of deformations of the full polyene chain in the total loss. The weights were chosen manually with the aim of reducing the loss terms for all deformation types and all molecules uniformly, without overfitting to any single contribution.

### Training

The initial parameters for the trained force field were imported from the GAFF2 force field. Atomic charges were assigned using the AM1-BCC method^36^, and Lennard-Jones parameters were kept at their GAFF2 values without optimization. Only the bonded parameters were trained. We adopted the GAFF2 atom typing scheme with one modification. In early experiments, we observed that training parameters for carotenoid end groups could conflict with parameters for sugars. To separate these parameter sets, sp³ carbons in six-membered carotenoid rings were assigned the atom type “c6,” while sp³ carbons in six-membered sugars were assigned the “c3” atom type. All other atom types followed the standard GAFF2 assignment.

In the GAFF2 polyene chain, single bonds are described by the atom-type pairs ce–ce or cf–cf, and double bonds by ce=cf (where ce and cf are inner sp^2^ carbons in conjugated chain systems). Swapping ce and cf yields a chemically equivalent but formally distinct atom-type assignment. To make the loss function symmetric with respect to this swap, the total loss was computed as the sum of losses evaluated for both typing variants. The analogous cg ↔ ch swap (where cg and ch are sp carbons in conjugated triple bonds) was not explicitly enforced. In the training and test sets, triple-bond-containing carotenoids analogous to alloxanthin were present, and in these molecules the ch atom was consistently positioned closer to the ring.

We optimized the following parameters: equilibrium bond lengths and their force constants, equilibrium angles and their force constants, and dihedral force constants. Phases and periodicities of dihedral terms were not optimized. GAFF2 assigns identical dihedral parameters to different atom-type quartets. At this training stage, in such cases, we optimized a single force constant shared across all equivalent quartets. The total number of GAFF2 parameters required to describe all molecules in this work exceeds 500. Training such a large number of parameters would likely lead to severe overfitting. We therefore selected a smaller subset of the most relevant parameters.

To identify these parameters, we computed the gradient of the loss function with respect to each parameter. For equilibrium bond lengths, the absolute values of the partial derivatives were sorted, and the 12 parameters with the largest magnitude were retained. For each selected bond, both the equilibrium length and the force constant were included in the optimization. The same procedure was applied to equilibrium angles: 13 angles were selected, and both the equilibrium angle and the force constant were optimized for each. Finally, 12 dihedral force constants with the largest gradient magnitudes were selected. The specific numbers of selected parameters (12 bonds, 13 angles, 12 dihedrals) were chosen manually by examining the relevance of the atom types. The total number of optimized parameters was thus 62 (obtained as 12×2+13×2+12).

Parameter optimization was performed using cyclic coordinate descent. Parameters were optimized in random order over three epochs. Each individual parameter was optimized via grid search, after which a cubic spline was fitted to the grid points to refine the optimal value.

After optimizing the initial 62 parameters, we further trained five additional dihedral parameters specific to the polyene backbone and carotenoid ring rotations. These were optimized in the same manner over four epochs. The total number of parameters optimized in this work was therefore 67. The complete set of 67 optimized parameters from both stages is provided as a Supplementary File 1.

Acylation with fatty acids is a common covalent modification of carotenoids, making the accurate description of long alkanes and alkenes critically important. In addition to training parameters for these molecules, we explored the possibility of adapting corresponding parameters from the Lipid21 force field. In Lipid21, dihedral energy profiles are fitted using a larger number of Fourier components than in GAFF2. To test whether adopting Lipid21 parameters improves simulation quality, we constructed a hybrid CARS+Lipid21 force field. In this hybrid scheme, alkane parameters are fully identical to those found in Lipid21. Alkene parameters differ from those in Lipid21 only slightly due to the Lennard-Jones parameters for sp² carbon and its bonded hydrogen being retained from GAFF2.

### Validation

Both the CARS and CARS+Lipid21 force fields substantially reduced the total loss compared to GAFF2, both across all molecules in the training set and within each molecular category (carotenoid end groups, lipids, sugars, etc.; Figure 2a). When the loss was decomposed by individual molecules and deformation methods, we observed an improvement in nearly all cases (Figure 2b). Only a few exceptions were found.

**Figure 2.**
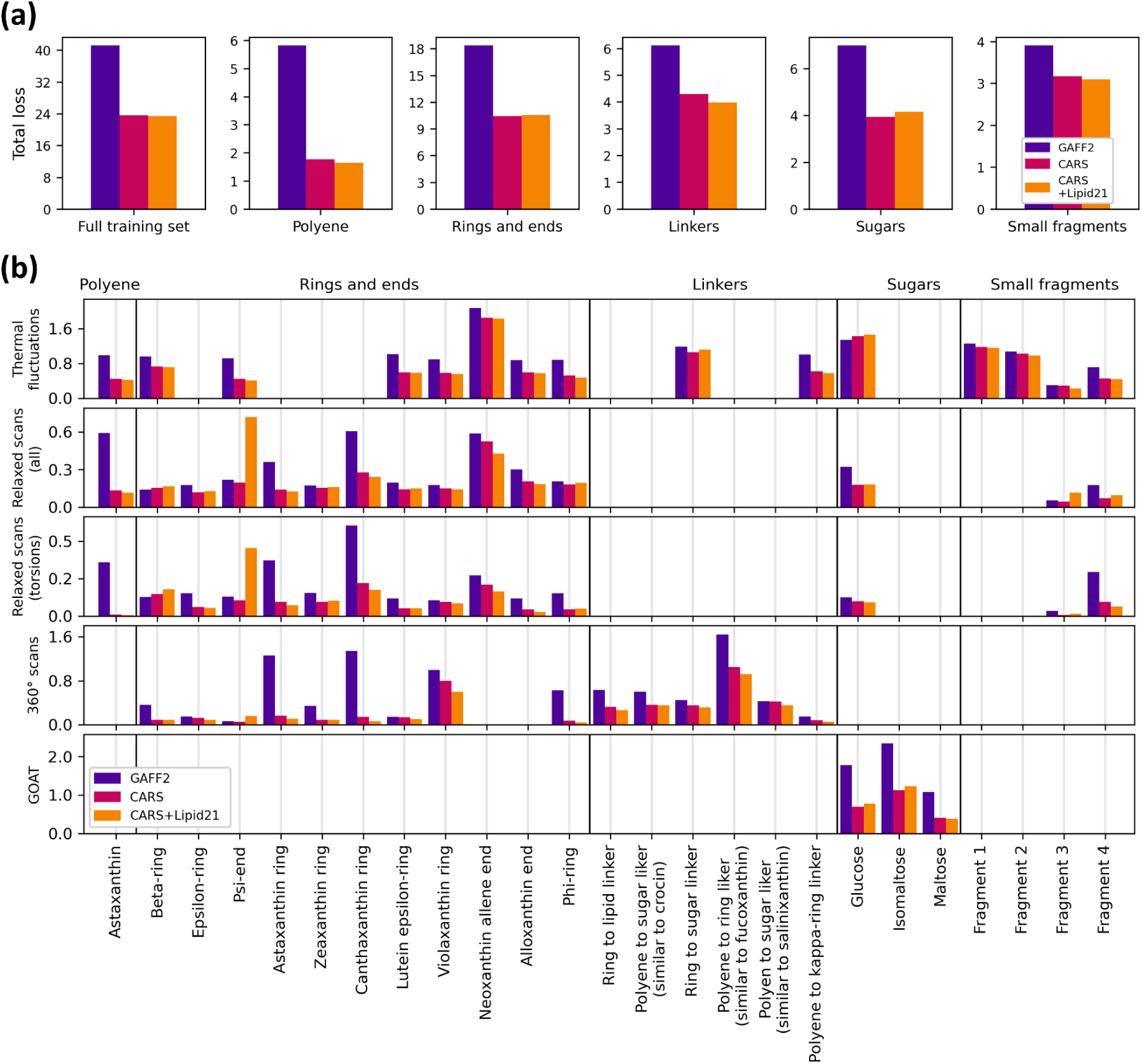
Evaluation of the developed force fields on the training set. (a) Total loss (MSE) for the GAFF2, CARS, and hybrid CARS+Lipid21 force fields, aggregated by molecular class. (b) Weighted loss contributions on the training set, decomposed by deformation protocol: molecular dynamics, relaxed scans (all internal coordinates, dihedral-only, and full 360° dihedral), and GOAT conformational sampling. For all deformation types except full 360° dihedral scans, the loss is reported as 1 − R^2^; for full 360° dihedral scans, the loss is MSE (scaled by 1 kcal²/mol²). The displayed losses are scaled to balance the contribution of each deformation type (see Results for weights). An additional factor of 3 was applied to all polyene contributions in the total loss calculation.

For the psi-end fragment, CARS+Lipid21 showed a markedly higher error than GAFF2 on relaxed surface scans. This is caused by the fact that the chemical environment of sp³ carbons in this molecule differs significantly from the sp³ environments present in the Lipid21 training set. In contrast, the pure CARS force field improved the predictions relative to GAFF2 for this fragment.

For glucose, both CARS and CARS+Lipid21 performed slightly worse than GAFF2 on the energies of MD-sampled conformations, but they yielded lower errors for GOAT-sampled conformers and relaxed surface scans. For the beta-ring, the trained force fields were slightly less accurate than GAFF2 on relaxed surface scans, whereas the energy profile for the full 360° rotation of the ring was improved.

To evaluate the developed force fields, CARS and CARS+Lipid21, on systems beyond the training set, we assembled a test set of 16 molecules encompassing the key structural motifs relevant to carotenoid chemistry (Figure 3). The carotenoid subset included beta-carotene as the simplest polyene, lutein with its epsilon- and beta-rings, alloxanthin with two triple bonds, neoxanthin containing an allenic bond, siphonoxanthin acylated with a fatty acid residue, crocin carrying two sugar moieties, bacterioruberin with an elongated polyene chain, oscillol-dirhamnoside combining an extended conjugation with glycosylation, and thermozeaxanthin bearing both a sugar and a lipid modification. The lipid subset comprised squalene, octadecane, and phytane, representing branched, linear, and polyunsaturated hydrocarbon chains. The sugar subset contained the monosaccharides glucose and rhamnose and the disaccharides lactose and sucrose. This selection covers the principal chemical features (glycosylation, acylation, extended conjugation, and different ring types) that the force fields must accurately describe. For each molecule in the test set, 200 conformations for carotenoids and lipids and 100 conformations for sugars were sampled from a 1 ns MD simulation in explicit TIP3P water, and the force-field energies were compared to QM reference energies using the MSE metric.

**Figure 3.**
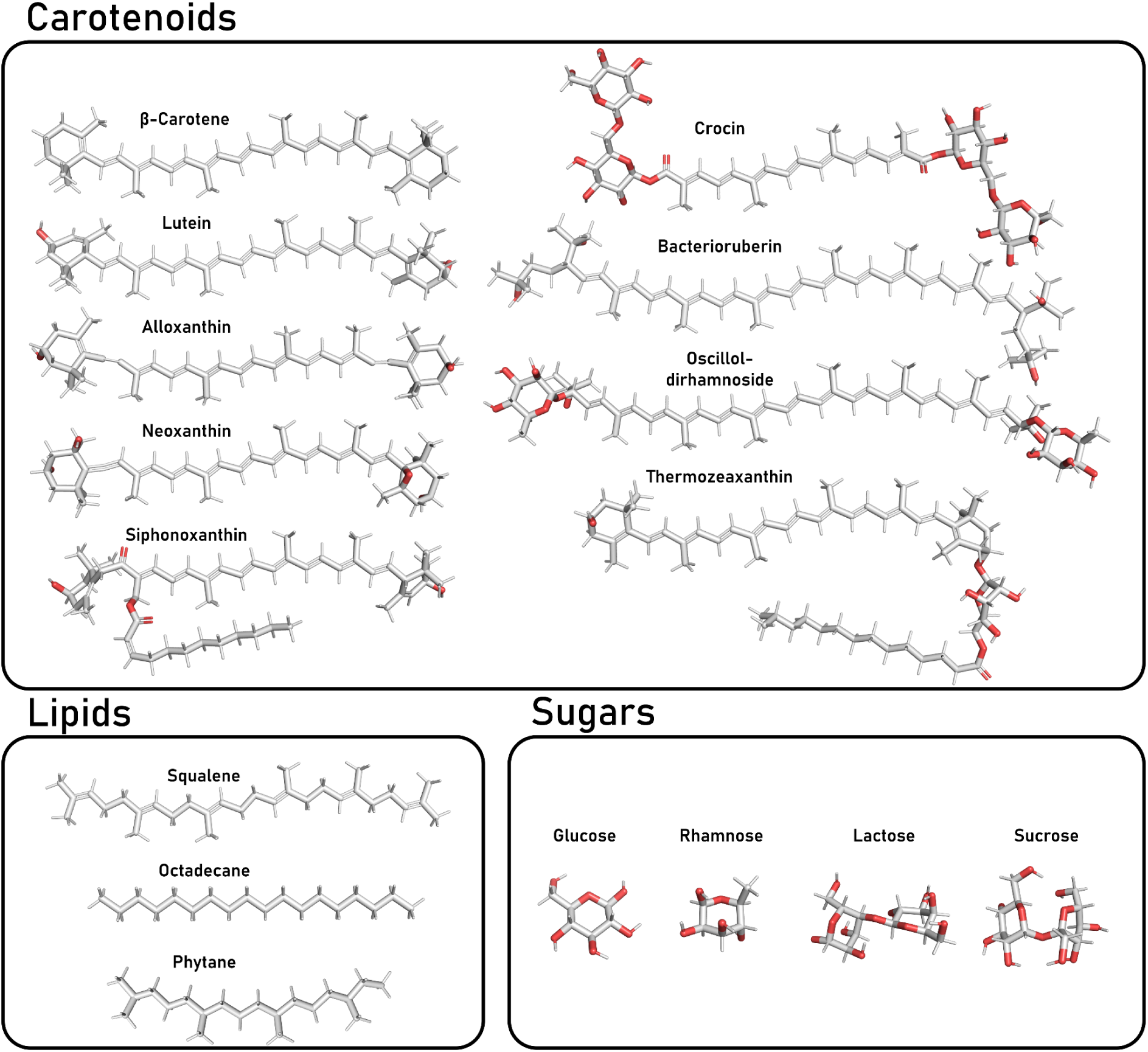
Test set for the CARS force field validation. All molecules were subjected to molecular dynamics simulations in explicit water at 300 K to generate conformational ensembles for subsequent energy evaluation.

For carotenoids, both CARS and CARS+Lipid21 showed substantially lower MSE values than GAFF2 across all nine molecules (Figure 4). For sugars, GAFF2 already outperformed the specialized GLYCAM_06j force field^37^ on all four sugar molecules. CARS, CARS+Lipid21, and GAFF2 exhibited comparable accuracy, with CARS achieving the lowest total MSE summed over the four sugar molecules. For lipids, GAFF2 gave the highest MSE among all tested force fields. CARS improved upon GAFF2 for squalene, octadecane, and phytane. CARS+Lipid21 performed nearly identically to the original Lipid21 force field and was more accurate than CARS alone, with the largest gains observed for squalene and octadecane.

**Figure 4.**
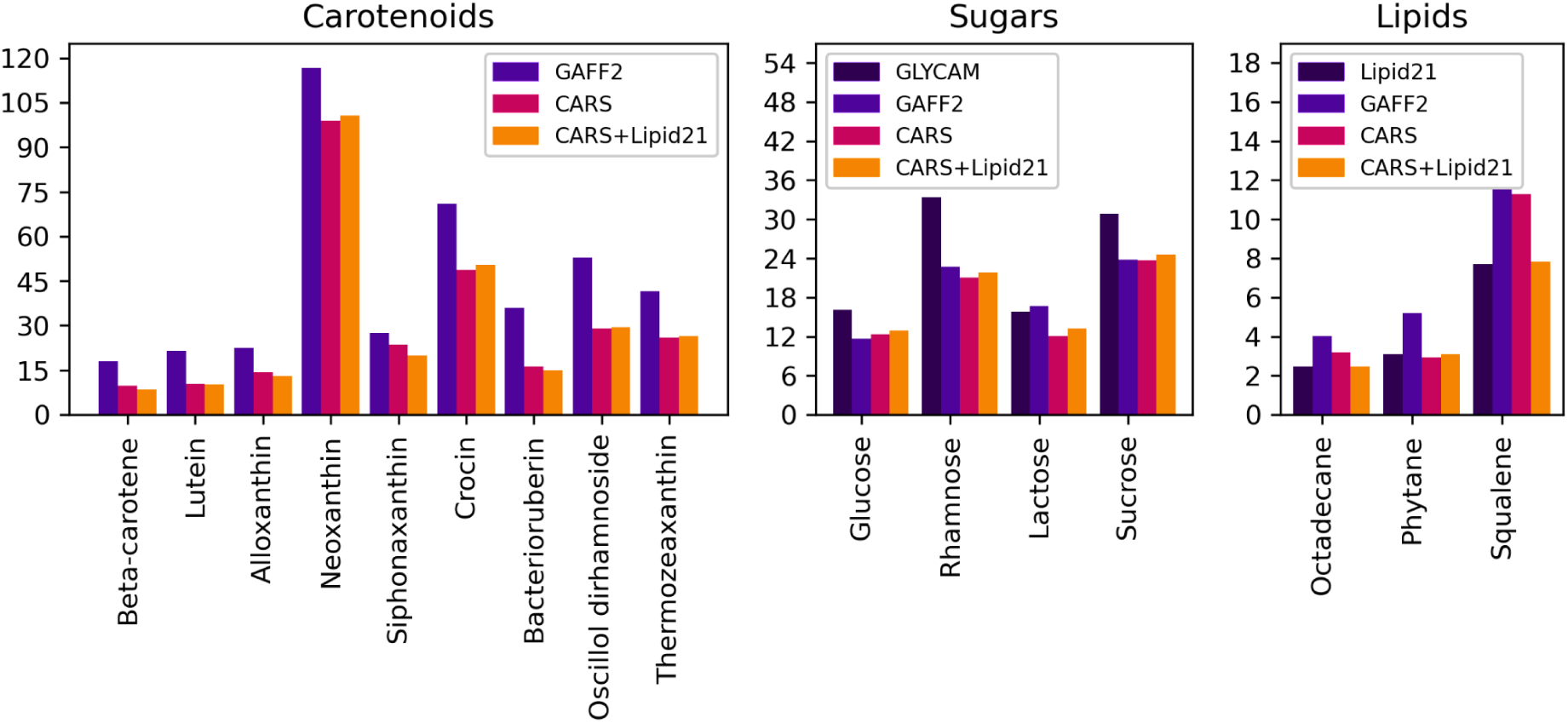
Evaluation of the force fields on the test set. Mean squared errors (MSE) for conformers sampled from MD simulations are shown without scaling. GLYCAM stands for GLYCAM_06j force field. Missing parameters in Lipid21 force filed for squalene are taken from GAFF2.

To assess the accuracy of geometry optimization, we selected two carotenoids: beta-carotene and oscillol-dirhamnoside, the latter combining a long polyene chain with sugar moieties. Reference geometries were optimized at the r2SCAN-3c level. The same molecules were then optimized with GAFF2, CARS, and CARS+Lipid21, and the RMSD values for all carbon atoms were computed relative to the QM reference (Figure 5). For beta-carotene, GAFF2 yielded an RMSD of 0.311 Å, CARS reduced it to 0.168 Å, and CARS+Lipid21 further lowered it to 0.107 Å. For the larger and more flexible oscillol-dirhamnoside, GAFF2 produced an RMSD of 0.844 Å, while CARS and CARS+Lipid21 achieved comparable values of 0.325 Å and 0.344 Å, respectively. Therefore, both trained force fields provide more accurate geometries than GAFF2.

**Figure 5.**
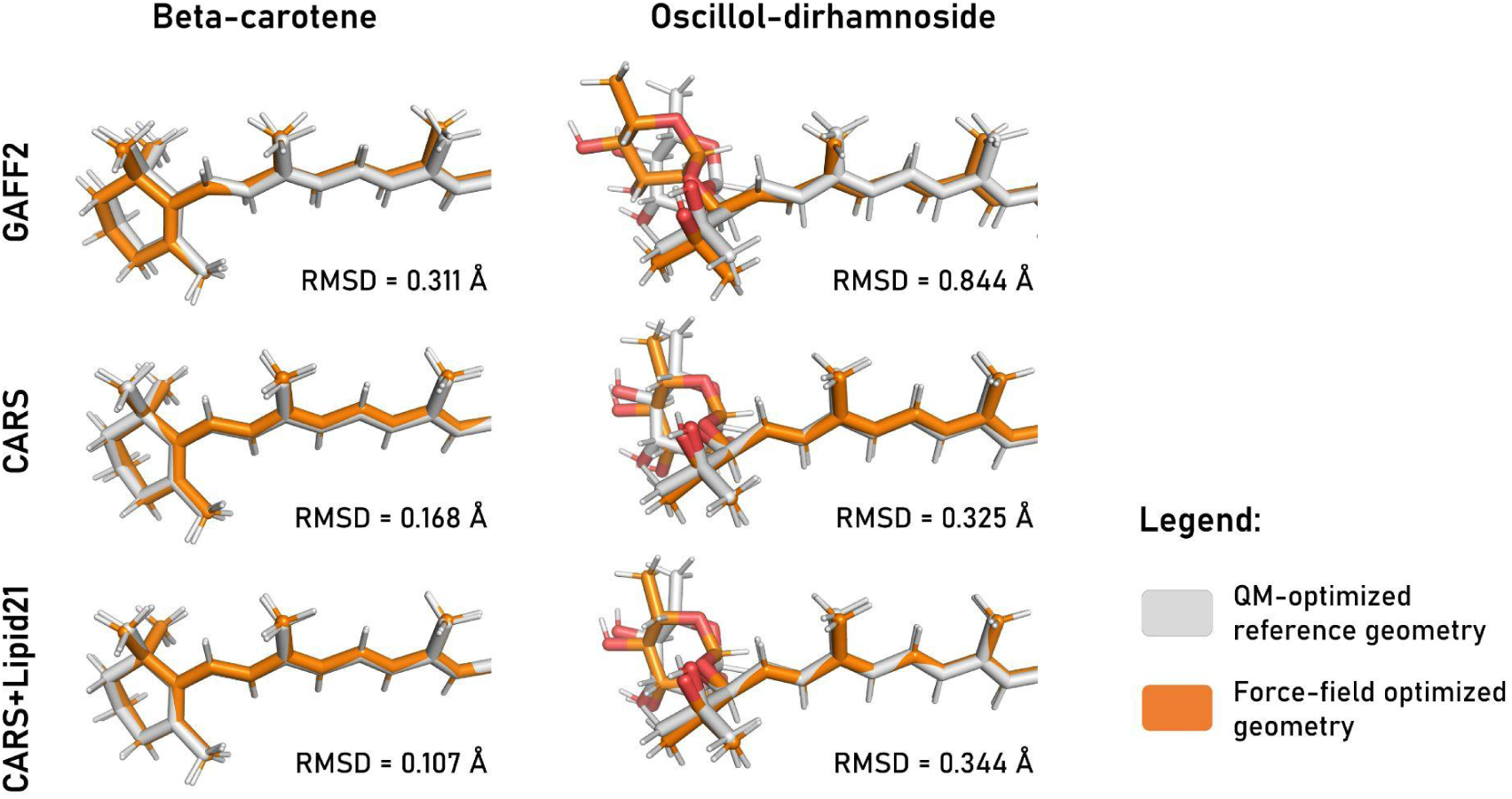
Comparison of optimized geometries for beta-carotene and oscillol-dirhamnoside using reference QM-method (shown in light gray) and GAFF2, CARS, and CARS+Lipid21 force fields (shown in orange). The figure shows six structural alignments (superimposed on carbon atoms) with RMSD values (Å) reported for each alignment.

Relaxed dihedral scans for carotenoid end groups rotations (Figure 6a) showed that GAFF2 already reproduces the reference r2SCAN-3c profiles reasonably well for the epsilon-ring, kappa-ring, psi-end, and neoxanthin ring torsions. In these cases, CARS and CARS+Lipid21 provided only marginal improvement, with CARS+Lipid21 slightly degrading the psi-end scans, consistent with the trend observed earlier. For the beta-ring and psi-ring torsions, however, GAFF2 exhibited large deviations from the reference, and both trained force fields substantially improved the agreement.

**Figure 6.**
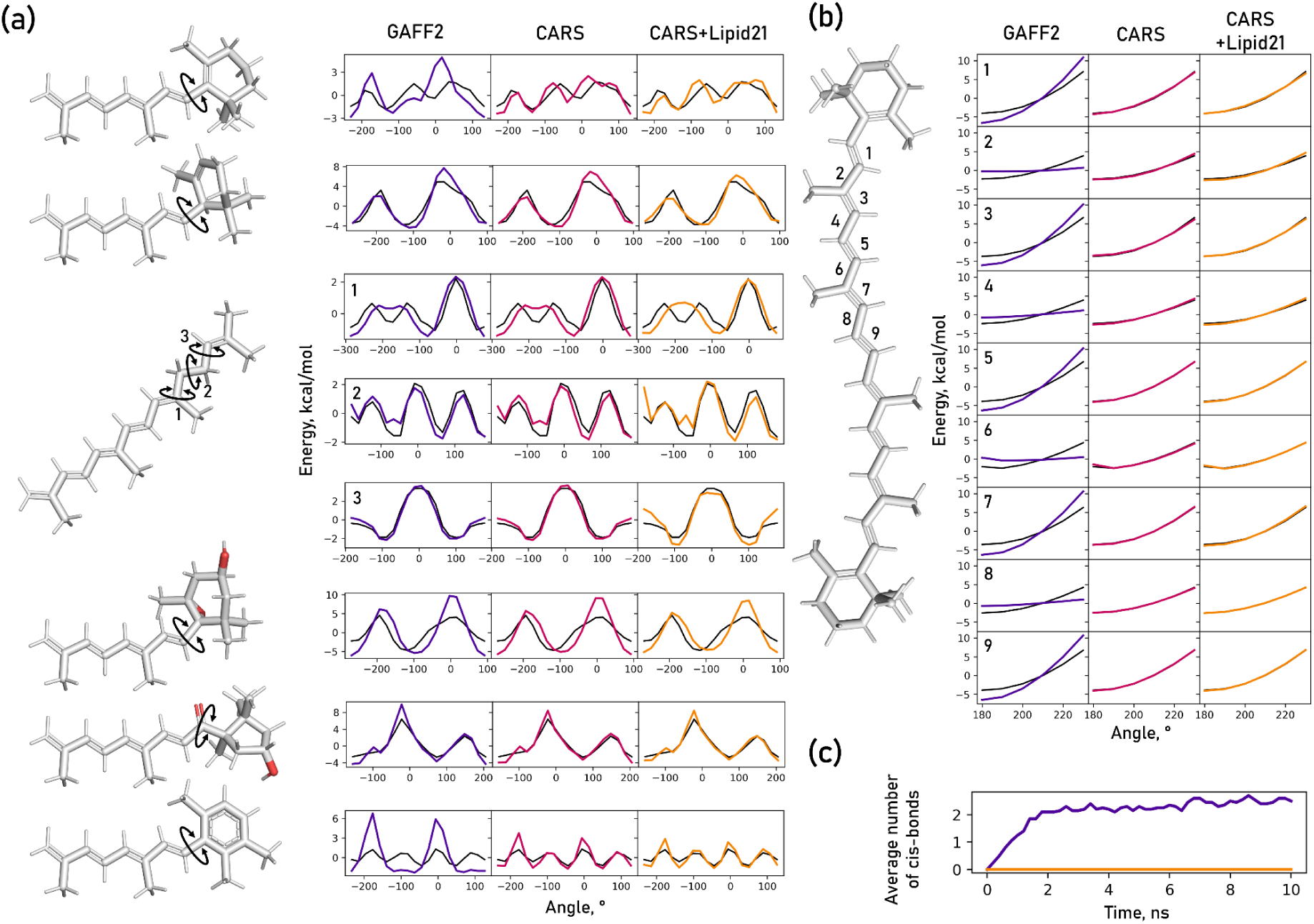
Validation of dihedral potentials. (a) 360° relaxed scans of the head group rotation in representative carotenoids, comparing GAFF2 (deep purple), CARS (ruby red), and CARS+Lipid21 (orange) against the QM reference (black). (b) Relaxed scans along selected polyene dihedral angles in beta-carotene, using the same colour scheme. (c) Time evolution of the average number of *cis* double bonds in beta-carotene during explicit-solvent MD simulations. Experimentally, cis–trans isomerization of beta-carotene does not occur on the nanosecond timescale^38^. GAFF2 shows a steady increase in *cis* isomer count over time, while CARS and CARS+Lipid21 maintain the expected all-*trans* population throughout the trajectory.

For polyene torsions, GAFF2 systematically underestimated the stiffness of single bonds and overestimated the stiffness of double bonds (Figure 6b). Because beta-carotene is experimentally known to keep its polyene chain conformation on the nanosecond timescale^38^, this imbalance constitutes a serious artifact. In 10 ns trajectories of beta-carotene in explicit water, the average number of *cis* bonds rose progressively with GAFF2, whereas both CARS and CARS+Lipid21 maintained the all-trans configuration throughout the simulations (Figure 6c).

To further benchmark the developed force fields, we considered zeaxanthin, a relatively simple symmetric carotenoid. We compared the performance of our models not only with general force fields (OPLS-AA, CGenFF, GAFF2), but also with the zeaxanthin-specific parameters from Prandi et al. (Figure 7).

**Figure 7.**
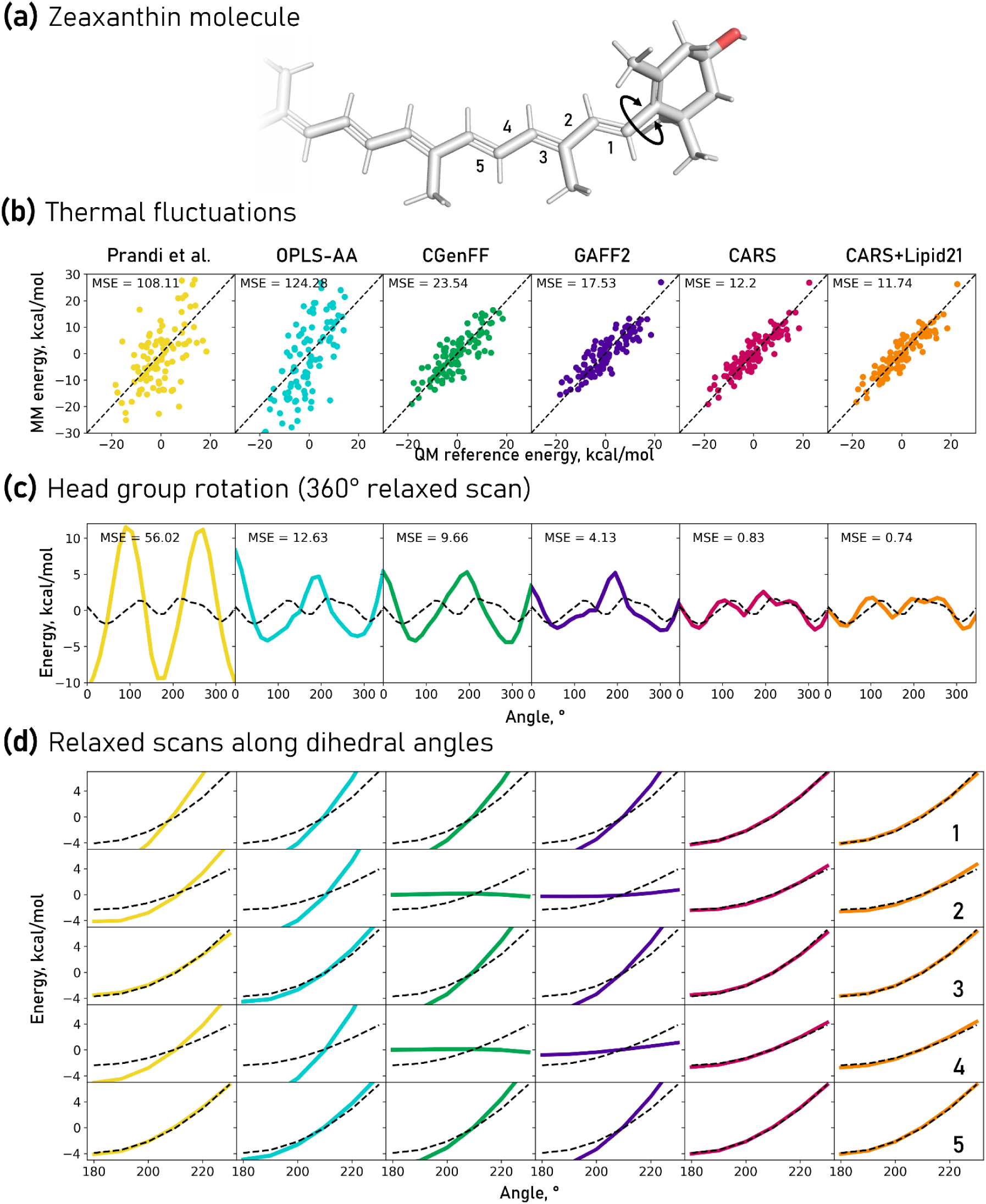
Evaluation of general force fields (OPLS-AA, CGenFF, GAFF2, CARS, and CARS+Lipid21) and the zeaxanthin-specific force field developed by Prandi et al^15^. (a) Structure of zeaxanthin showing the head-group rotation (double arrow) and the numbering of polyene bonds used in panel (d). (b) Scatter plots of QM versus MM energies for conformers sampled from one MD trajectory. (c) 360° relaxed scans of the head-group dihedral: QM reference (black) and force-field predictions (coloured). (d) Relaxed scans of the polyene dihedral angles corresponding to the bonds indicated in (a).

For the set of MD-sampled conformers, CGenFF, GAFF2, CARS, and CARS+Lipid21 all reproduced the reference QM energies with low errors (Figure 7b). OPLS-AA and the Prandi parameters gave substantially larger deviations, with MSE values exceeding 100 kcal² mol⁻². CARS+Lipid21 achieved the lowest MSE among all tested models. For the 360° rotation of the beta-ring head group (Figure 7c), the Prandi parameters showed the largest discrepancy (MSE > 50 kcal² mol⁻²), while CARS+Lipid21 provided the closest match to the reference. OPLS-AA, CGenFF, and GAFF2 fell in between, with GAFF2 being the most accurate of the three. Relaxed scans along the polyene chain (Figure 7d) highlighted systematic differences: OPLS-AA and the Prandi parameters overestimated the torsional stiffness, CGenFF and GAFF2 underestimated the stiffness of single bonds, whereas CARS and CARS+Lipid21 gave the best agreement with the QM profiles.

However, benchmark energies alone do not guarantee correct dynamics: the lower polyene dihedral force constants in CARS, compared to the stiffer torsions in OPLS-AA or the Prandi parameters, could in principle allow non-physical isomerization. Additionally, the trained parameter set must permit numerically stable simulation at a 2 fs integration step. To assess whether the improved energy profiles of CARS+Lipid21 translate into stable and physically plausible MD trajectories, we performed microsecond-scale simulations of three carotenoid–protein complexes in explicit solvent.

**Figure 8.**
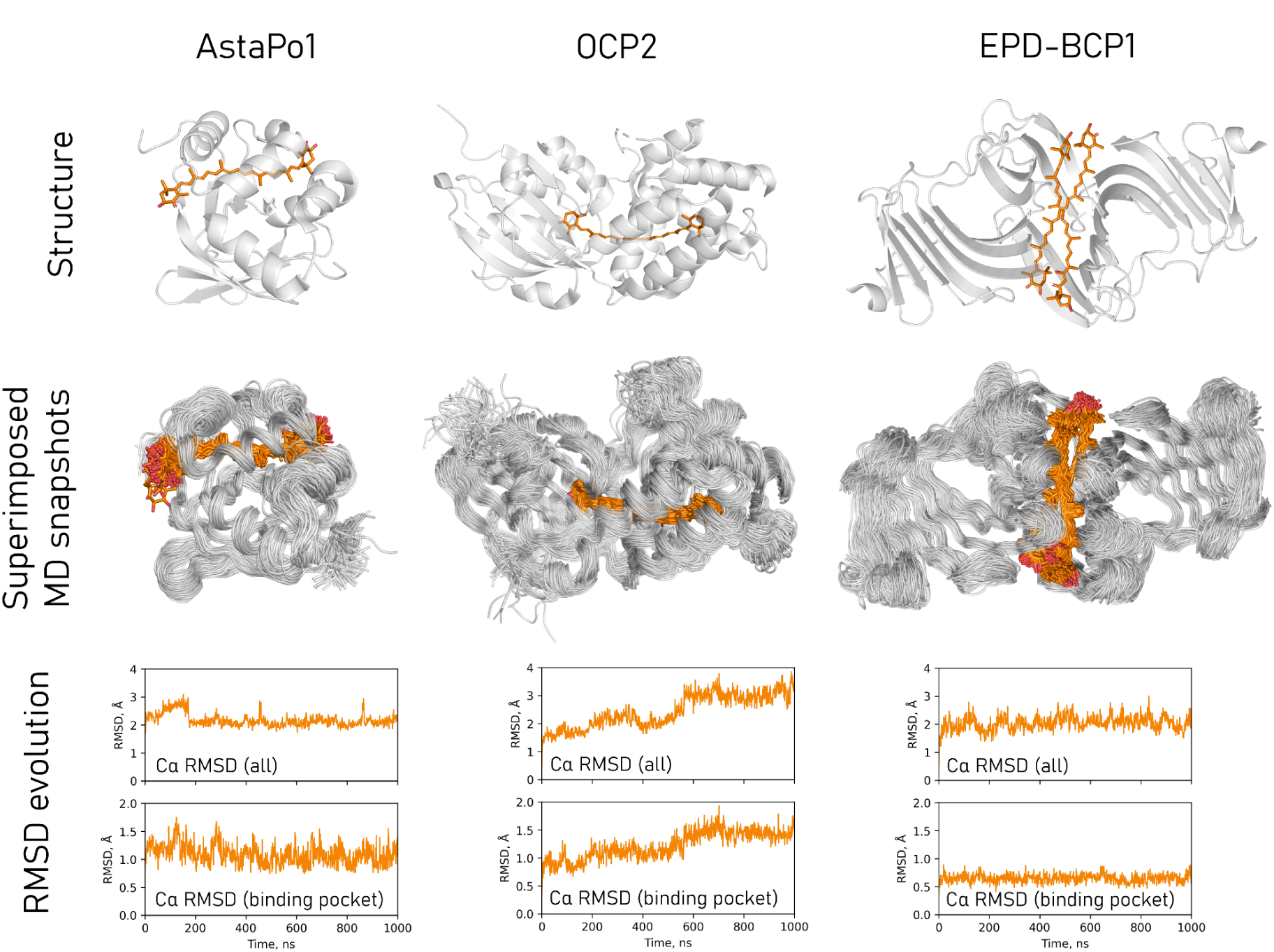
Evaluation of CARS in MD simulations of soluble proteins. (Top row) Structures of the three carotenoid-binding proteins: AstaPo1 bound to astaxanthin (PDB ID 8C18^39^), OCP2 bound to echinenone (PDB ID 8PYH^40^), and EPB-BCP1 bound to astaxanthin and mytiloxanthin (PDB ID 8I34^41^). (Middle row) Superimposed snapshots from 1 μs molecular dynamics simulations in water. (Bottom row) Cα RMSD as a function of time, shown for all residues and for the binding pocket. The pocket is defined as all residues having the Cα atom within 7 Å of the chromophore in the initial structure. All simulations were performed with the CARS+Lipid21 force field for carotenoids and ff14SB for proteins.

We selected three soluble proteins: AstaPo1 in complex with astaxanthin (PDB ID 8C18^39^), OCP2 with echinenone (PDB ID 8PYH^40^), and the blue carotenoid glycoprotein EPB-BCP1, which binds both astaxanthin and mytiloxanthin (PDB ID 8I34^41^). All systems were simulated with the CARS+Lipid21 force field for 1 μs in explicit water. No *cis*-isomerization of the carotenoids was observed in any of the trajectories, and the simulations ran without numerical instability at a 2 fs time step. In AstaPo1 and EPB-BCP1, the Cα RMSD of the binding pocket remained low throughout the trajectory (approximately 1.2 Å and 0.6 Å, respectively). In OCP2, the overall protein RMSD increased steadily and reached nearly 4 Å by the end of the simulation, while the pocket RMSD rose to about 1.5 Å. This drift likely reflects genuine conformational flexibility of OCP2. Overall, the simulations confirm that CARS+Lipid21 maintains polyene geometry, numerical stability, and pocket integrity on the microsecond timescale.

Many carotenoids function in lipid membranes and interact with membrane proteins. To test the behaviour of carotenoids modeled with CARS+Lipid21 in a membrane environment, we simulated two carotenoid-containing rhodopsins: the myxol-bound proteorhodopsin NM-R1 (PDB ID 9JOV^42^) and xanthorhodopsin with salinixanthin (PDB ID 3DDL^43^). In both systems, the carotenoid lies at the protein surface, with one endgroup contacting the retinal and the other, often less ordered, extending toward the lipid headgroups.

Both systems were simulated for 1 μs in an explicit lipid bilayer. The trajectories showed no numerical instability and no isomerization of the carotenoids. In the crystal structures, the B-factors reflecting thermal fluctuations of the carotenoids are low near the retinal-proximal endgroup and high at the opposite end; for salinixanthin, the sugar and lipid substituents also display elevated B-factors. The simulations closely reproduce this pattern: the endgroup closest to the retinal stays tightly anchored to the protein, while the distal end remains highly dynamic. The qualitative agreement between the crystallographic B-factors and the simulated flexibility suggests that the force field preserves the expected partitioning of rigidity and disorder in membrane-bound carotenoids.

**Figure 9.**
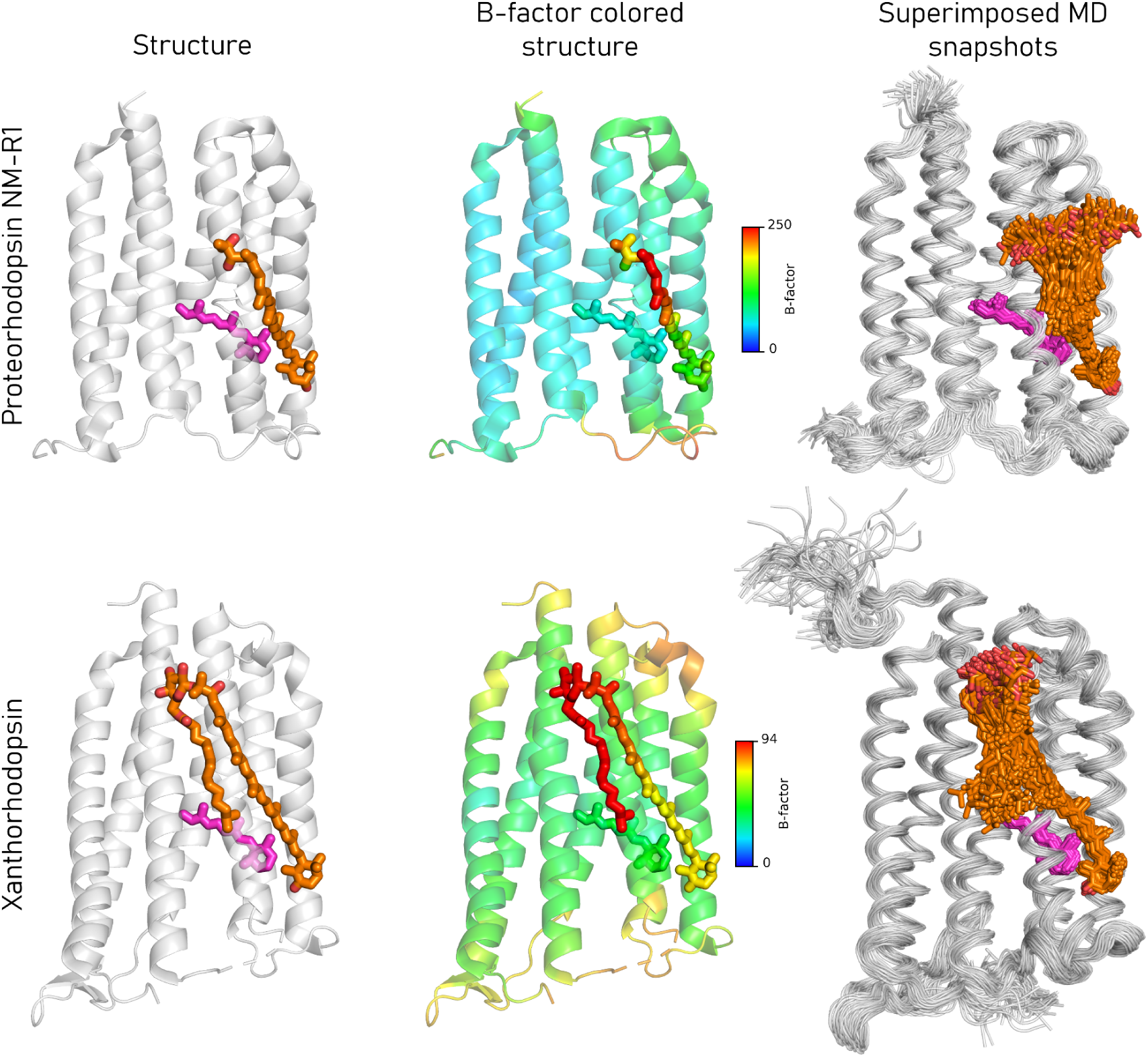
Evaluation of CARS in MD simulations of membrane proteins. (Left column) Structures of proteorhodopsin NM-R1 bound to myxol (PDB ID 9JOV^42^) and xanthorhodopsin bound to salinixanthin (PDB ID 3DDL^43^). The carotenoids are shown in orange and the retinal is in magenta. (Middle column) The same structures colored by crystallographic B-factor with a rainbow gradient. While the proteins and retinal-proximal groups of carotenoids are relatively ordered, the retinal-distant parts of carotenoids are strongly disordered. (Right column) Superimposed snapshots from 1 μs molecular dynamics simulations in a lipid membrane. The pattern of carotenoid fluctuations closely matches the pattern of B factor distribution in experimental structures. All simulations were performed with the CARS+Lipid21 force field for carotenoids, Lipid21 for lipids and ff14SB for proteins.

The CARS and CARS+Lipid21 parameter sets may also improve the description of other families of biomolecules. One example is glycolipids, which are abundant in thylakoid membranes, primarily as monogalactosyldiacylglycerol (MGDG) and digalactosyldiacylglycerol (DGDG); neither is covered by Lipid21 alone. Because CARS+Lipid21 inherits lipid-tail parameters from Lipid21 and includes sugar moieties in its training data, its accuracy on glycolipids is of practical interest.

We examined two test molecules: tributyrin, a simple triacylglycerol, was chosen to assess the description of the glycerol ester backbone, and MGDG(4:0/4:0) was selected as the simple MGDG representative (Figure S1). For tributyrin, CARS+Lipid21 reduced the MSE on GOAT-sampled conformers from 7.4 (kcal/mol)² (GAFF2) to 1.7 (kcal/mol)², close to the Lipid21 value of 1.6 (kcal/mol)². On MD-sampled conformers, the MSE decreased from 18.5 to 15.5 (kcal/mol)² (Lipid21: 9.6 (kcal/mol)²). For MGDG(4:0/4:0), the improvement on GOAT conformers was more pronounced: the MSE dropped from 27.2 (kcal/mol)² (GAFF2) to 3.7 (kcal/mol)². On MD-sampled conformers, the MSE fell from 27.0 to 23.7 (kcal/mol)². In all cases, CARS+Lipid21 outperformed GAFF2.

Another example is polyene macrolide antifungals. These molecules consist of a long conjugated polyene chain and sugar residues, a combination that closely matches the fragments present in the CARS training set. We tested CARS on amphotericin B (Figure S2). For MD-sampled conformers, CARS reduced the MSE from 74.9 (kcal/mol)² (GAFF2) to 65.3 (kcal/mol)². Given the consistent polyene stability observed in all CARS simulations of carotenoids, the same parameter set is expected to maintain the all-*trans* configuration in this class of molecules as well.

## Discussion

Carotenoids are frequently simulated with classical force fields, but no existing parameter set describes their full structural diversity in a unified way. Here, we introduced CARS, a modular force field designed to cover most known carotenoid species within a single parameter framework.

CARS accurately reproduces the torsional energy profiles of the polyene backbone and of the head-group rotations. Equilibrium geometries improve relative to GAFF2. For sugars, CARS surpasses GLYCAM_06j in the energies of thermal deformations and significantly improves conformational energies compared to GAFF2. On zeaxanthin, CARS outperforms OPLS-AA^8^, CGenFF^44^, GAFF2^6^, and parameters developed by Prandi et al.^15^ across all three validation metrics: conformer energies, ring rotation, and polyene scans. The relative performance of CARS and CARS+Lipid21 depends on the chemical composition of the carotenoid: for molecules that carry a saturated or nearly saturated lipid tail, CARS+Lipid21 is the preferred choice, inheriting the accuracy of Lipid21 for the lipid moiety without degrading the polyene; for carotenoids that contain sugars but no lipid modification, the pure CARS force field gives slightly more accurate results, and it is clearly preferable when the molecule includes a psi-end group.

CARS was trained on a set of fragments covering the most common carotenoid end-groups, sugars, and lipid attachments. For any molecule whose chemical groups are represented in that set, the optimised bonded parameters are expected to work as validated; chemical groups absent from the training data will be described with GAFF2 accuracy. This modular design makes CARS applicable beyond carotenoids, to other natural products that share the same building blocks. One example is polyene antimycotics such as amphotericin B, whose heptaene chain and sugar residue closely match the fragments in the training set. We showed that CARS improves conformational energies of amphotericin B relative to GAFF2, providing a basis for simulating its interaction with fungal membranes^45^.

The training set leaves out several structural motifs found in natural carotenoids. It does not include end-groups that contain two fused cycles, such as those in mutatochrome and cycloviolaxanthin, nor does it cover molecules in which a cycle is inserted in the middle of the polyene chain as in peridinin. Charged carotenoids, in particular apocarotenoids that carry a free carboxyl group, were also not part of the training data, so their behaviour when modeled using CARS remains to be tested.

Atomic charges for all molecules in the training set were assigned using the AM1-BCC method^36^. This scheme is computationally inexpensive, widely available, and has a long history of use together with GAFF and GAFF2. Several new fast charge assignment schemes, such as ABCG2^46^ and EspalomaCharge^47^, have recently become available and their performance with CARS remains to be evaluated. The more rigorous protocol adopted in Lipid21^4^ (RESP charges^48^ averaged over multiple MD-derived conformations) remains an open option, but it is prohibitively expensive because of the large number of atoms in typical carotenoids. Lennard-Jones parameters were taken from GAFF2 without adjustment; refining them for the sp² carbons of the polyene chain against experimental data could further improve intermolecular packing and is left for future work.

Several areas could directly benefit from the improved accuracy that is provided by CARS, with one of them being the simulation of photosynthetic systems. Carotenoids are essential components of photosystems^49^ and light-harvesting complexes (LHCs)^50^. In these systems, carotenoids act as light-harvesting pigments and as photoprotective agents that quench singlet oxygen^51^. Reliable simulations of these proteins therefore require parameters that accurately describe the bound carotenoids. A further challenge for modeling photosynthetic membranes is that the major thylakoid glycolipids MGDG and DGDG^52^ are not parameterized in Lipid21. CARS+Lipid21 combines the accurate lipid-tail parameters of Lipid21 with the improved sugar and glycerol ester description of CARS relative to GAFF2. This combination provides a physically motivated starting point for building parameters for MGDG, DGDG, and other glycolipids in the future.

Beyond photosynthesis, carotenoid binding protein complexes are increasingly recognized as the molecular basis of coloration in animals. The blue hue of the lobster shell^53^, the bright yellow of gregarious locusts^54^, the striking blue of a marine sponge^41^, and the leaf-green camouflage of bush crickets^55^ all arise from specific protein scaffolds that bind carotenoids and tune their absorption. Reliable force fields for the carotenoid component are essential for understanding, through MD simulations combined with QM calculations, how the protein environment modulates the spectral properties of the pigment.

Microbial rhodopsins were recently highlighted as a large family of carotenoid-binding photoactive proteins^56^. Once thought to be a curious exception in *Salinibacter ruber*^43^, rhodopsin-carotenoid complexes are now found in diverse marine bacteria, suggesting a widespread strategy for solar energy capture. These proteins use a carotenoid as an accessory antenna that absorbs in the wavelength range not captured by the retinal chromophore, transfers excitation energy to it and thus extends the action spectrum. In these systems, the carotenoid resides at the protein–membrane interface, with one endgroup contacting the retinal and the other extending into the lipid bilayer. How the protein selects a specific carotenoid and how the chemical identity and dynamics of the carotenoid influence the efficiency of light harvesting and energy transfer to retinal are important questions^57^ that MD simulations with a reliable force field can help to address.

Overall, CARS and CARS+Lipid21 together provide a single parameter set for simulations of chemically diverse carotenoids, removing the need to develop individual parameters for each new molecule. These two force fields cover the most common end-groups, glycosylation, and lipid acylation types and are as simple to use as GAFF2. We anticipate that CARS will facilitate routine atomistic simulations of carotenoids across diverse biological systems and accelerate mechanistic studies of their structure, dynamics, and molecular recognition.

## Methods

### QM calculations

All quantum mechanical calculations were performed with ORCA 6.1.1^58^. Single-point energies were computed at the r2SCAN-3c^28^ level with the TightSCF convergence criterion. Reference geometries for beta-carotene and oscillol-dirhamnoside were optimized at the same r2SCAN-3c level.

Relaxed surface scans of internal coordinates were performed at the GFN2-xTB^32^ level using ORCA with default settings. Conformational sampling of sugars was carried out with the GOAT algorithm as implemented in ORCA, also at the GFN2-xTB level with default parameters. For all GOAT-sampled conformers, single-point energies were subsequently recalculated at the r2SCAN-3c level. For glucose, the local minima identified by GOAT were additionally reoptimized at the r2SCAN-3c level; for the remaining sugars, only single-point recalculations were performed.

### Molecule selection and initial geometry preparation

Molecular fragments for the training set were selected based on structures from the Carotenoids Database^2^ (accessed October 2025). The fragments were constructed manually in Avogadro 1.2.0^59^ and geometry-optimized using the GAFF force field as implemented in Avogadro. Molecules for the test set were obtained from PubChem^60^ as SMILES strings, built in Avogadro 1.2.0, and geometry-optimized with the same GAFF implementation. The complete lists of fragments and molecules included in the training and test sets are shown in Figures 1 and 3.

### Force field versions and parameter generation

GAFF2 (Version 2.2.20, March 2021) from AmberTools23^61^ was used as the baseline force field. GLYCAM_06j^37^ parameters and partial charges were obtained via the GLYCAM Web server^62^. OPLS-AA^8^ parameters and charges (1.14*CM1A) were generated with LigParGen^63^. CGenFF parameters and charges were obtained from the CGenFF server^44^. For all server-generated force fields, default settings were used.

### MD-based conformational sampling

Each molecule was solvated in a cubic box of TIP3P^34^ water with a minimum distance of 10 Å between the solute and the box edge. Systems were assembled using tleap from AmberTools23^61^. GAFF2 parameters and AM1-BCC charges were assigned to all molecules. Simulations were performed with OpenMM 8.2^64^ using a 2 fs time step, a Langevin integrator with a friction coefficient of 1 ps⁻¹, and a Monte Carlo barostat set to 1 atm. Each system was equilibrated for 10 ps in the NPT ensemble, followed by a 1 ns production NPT simulation, from which 1000 frames were extracted.

For carotenoid fragments and full carotenoids, unmodified GAFF2 leads to rapid *cis–trans* isomerization around single bonds of the polyene chain. To suppress this non-physical behavior during sampling, selected dihedral force constants were manually adjusted prior to the simulations. For fragments containing a half-polyene chain attached to a ring, the force constants for single bonds (atom-type quartets X-ce-ce-X and X-cf-cf-X) were increased from 1.00 to 1.23856 kcal/mol, and those for double bonds (X-ce-cf-X, X-cf-c2-X, X-ce-c2-X, X-c2-c2-X) were reduced from 6.65 to 3.89653 kcal/mol. For full carotenoids with an intact polyene chain, the corresponding values were set to 1.70 and 3.89653 kcal/mol, respectively. These empirically adjusted parameters allowed physically plausible sampling of the all-*trans* configuration. The final trained values from the first stage of CARS force field training are 1.71293 kcal/mol for single bonds and 4.48572 kcal/mol for double bonds.

### Calculation of the loss function

Atomic types and AM1-BCC charges were assigned to all molecules using antechamber from AmberTools23. The atom typing for the conjugated polyene chain (ce and cf types) was manually inspected and corrected where necessary to ensure a consistent assignment. Missing bonded parameters were generated with parmchk2 from AmberTools23 by analogy to existing GAFF2 parameters and stored as .frcmod files. A single topology file (.prmtop) containing all molecules of the training set was then assembled in tleap from AmberTools23, thereby defining the minimal parameter set required to describe the entire dataset. This unified parameter set could be modified programmatically using ParmEd^65^. After modifications, the .prmtop was split into individual topologies for each molecule, again using ParmEd. For each molecule, the corresponding topology and the pre-generated XYZ conformations were loaded into OpenMM to compute the MM energy at the GAFF2 level with the applied parameter changes. The total loss was then evaluated as a weighted sum of MSE and 1–R² contributions, partitioned by molecule and deformation type as described in the Results section.

MSE loss was defined as

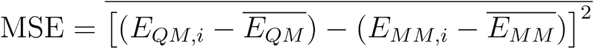

where E_QM,i_ and E_MM,i_ are the reference QM energy and the force-field energy of conformation i, respectively, and *E̅_QM_* and *E̅_MM_* are the mean values over all conformations. The overline on the outer bracket denotes averaging over all conformations.

The 1 − R^2^ loss was defined as

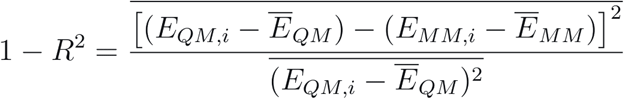

where E_QM,i_ and E_MM,i_ are the reference QM energy and the force-field energy of conformation i, respectively, *E̅_QM_* and *E̅_MM_* are the corresponding mean values over all conformations, and the overline denotes averaging over all conformations.

### Parameter optimization

Gradients of the loss function with respect to individual parameters were computed numerically by finite differences, f(x+Δx)−f(x); Δx=x/1000. The absolute gradient values were calculated for all bonded parameters in the unified topology. The 12 bond parameters, 13 angle parameters, and 12 dihedral force constants with the largest absolute gradients were selected for optimization. For each selected bond and angle, both the equilibrium value and the force constant were included.

Parameter optimization was performed by cyclic coordinate descent. A random permutation of the selected parameters was generated once and reused for all three epochs of the first training stage. For each parameter, a one-dimensional grid search was conducted around the current value. The search ranges were set to ±2% of the current value for equilibrium bond lengths and angles, ±10% for bond and angle force constants, and ±30% for dihedral force constants. Each grid contained seven equally spaced points. A cubic spline was fitted to the grid points, and the minimum of the spline was taken as the refined parameter value.

The first stage of training optimized 62 parameters over three epochs. In the second stage, five additional dihedral parameters specific to the polyene backbone and carotenoid ring rotations were added and optimized over four epochs, bringing the total number of trained parameters to 67. A fixed number of epochs was used in both stages as a safeguard against overfitting. The complete procedure was implemented in Python using ParmEd and OpenMM. The final parameter set is provided as Supplementary File 1.

### CARS+Lipid21 hybrid force field

The hybrid CARS+Lipid21 force field was constructed by incorporating the bonded parameters of Lipid21^4^ for saturated and unsaturated hydrocarbon chains into the CARS parameter set. First, the molecules squalene, phytane, and octadecane were assigned GAFF2 atom types. These types were then temporarily converted to the corresponding Lipid21 types (c3 → cD, c2 → cB, hc → hL, ha → hB) . A topology file (.prmtop) was built for the system containing all three molecules with Lipid21, and all parameters present in this file were extracted into an .frcmod file using ParmEd. The atom types in the .frcmod file were subsequently converted back to the original GAFF2 types, and the non-bonded parameters for atom types ha and c2 were discarded. The resulting.frcmod file was loaded into tleap after the CARS parameters, thereby replacing the corresponding GAFF2 bonded parameters for alkanes and alkenes with those of Lipid21.

#### Test set and validation methodology

A test set of 16 molecules (9 carotenoids, 3 lipids, 4 sugars; see Results for the full list) was used for independent assessment. Test set conformations were generated via MD simulations identical to those described in “MD-based conformational sampling”, with 200 frames extracted per each carotenoid and lipid, and 100 frames per each sugar. Reference energies for all test conformers were computed at the r2SCAN-3c level (TightSCF, ORCA 6.1.1).

Relaxed dihedral scans (both limited-range and 360°) used for validation were performed with GFN2-xTB geometry optimization and subsequent r2SCAN-3c single-point energy evaluation, exactly as for the training set. Geometry optimizations with GAFF2, CARS, and CARS+Lipid21 were carried out in ORCA 6.1.1 using the default optimizer. RMSD values for the carbon atoms were calculated in PyMOL with the align command (cycles=0).

To assess polyene stiffness, 20 independent 10 ns NPT MD simulations of beta-carotene were run under the conditions of “MD-based conformational sampling”, and the average number of *cis* bonds was monitored.

For comparison with published force fields, the zeaxanthin parameters of Prandi et al. were used in their original Amber format. OPLS-AA, CGenFF, and GAFF2 parameters were generated as described above. All force fields were evaluated on the same MD-sampled conformers and dihedral scans of zeaxanthin, using r2SCAN-3c as the reference.

#### MD systems assembly

Initial configurations were generated using tleap from AmberTools25 for systems of soluble proteins (PDB IDs 8C18, 8I34, 8PYH), and with packmol-memgen of AmberTools25 for microbial rhodopsin systems (PDB IDs 3DDL, 9JOV). Initial box sizes for 8C18, 8I34, and 8PYH were determined so that the minimum distance from the protein to the box edge was 2.0, 1.2, and 1.2 nm, respectively (using the SolvateBox command of tleap). The initial box size of rhodopsin systems was 10.0 × 10.0 × 10.6 nm. The membranes of rhodopsin systems consisted of POPE and POPG in a 3:1 ratio.

Systems were built with tleap from AmberTools25. ff14SB^66^ and TIP3P were used for proteins and water, respectively. Protonation states of titratable residues were assigned for pH 8 based on pKa values determined with PropKa3^67^. The only exception was the carboxyl counterion of the retinal Schiff base in rhodopsins, which was kept deprotonated. POPE and POPG were modelled with Lipid21. Retinal and carotenoids were modelled with CARS+Lipid21; the boundary between the ff14SB and CARS+Lipid21 parameter sets was placed at the midpoint of the lysine side chain that forms the Schiff base linkage to retinal. Partial charges for carotenoids were obtained by geometry optimization at the GFN2-xTB level, followed by a single-point HF/6-31G*^68^ calculation and RESP^48^ fitting with MultiWFN^69^. Protein glycosylation sites and sugars were described with GLYCAM_06j.

The 8I34 system was converted to GROMACS file format using acpype^70^ because GLYCAM_06j parameters were not directly convertible via ParmEd. All other systems were converted with ParmEd.

#### Simulation parameters

All simulations were performed with GROMACS 2024.5^71^ using a time step of 2 fs. The Verlet cutoff scheme was used. The neighbor list update interval was determined adaptively using default GROMACS parameters. Lennard-Jones interactions were truncated at a cutoff of 9 Å, with long-range dispersion corrections applied to energy and pressure. The PME^72^ scheme was used for Coulomb interactions with adaptive parameters based on default GROMACS settings.

Temperature was controlled with the V-rescale thermostat^73^ at 300 K with a coupling time constant τ = 1 ps. Pressure was controlled with the C-rescale barostat^74^ at 1 bar with a coupling time constant τ = 5 ps and a compressibility of 4.5 × 10⁻⁵ bar⁻¹. For soluble protein systems, an isotropic barostat was used and two temperature-coupling groups were defined: protein–ligand and solvent. For membrane protein systems, a semi-isotropic barostat with equal parameters along the z-axis and the xy-plane was used, and three temperature-coupling groups were defined: protein–ligand, solvent, and lipids. Only the linear velocity of the center of mass was removed.

All systems were energy-minimized (5000 steps) and subsequently equilibrated in the NVT and NPT ensembles before production runs. NVT equilibration was performed for 100 ps with a 1 fs time step for soluble protein systems and for 100 ns with a 1 fs time step for membrane systems. Thermostat and barostat parameters were the same as described above. All heavy atoms of the protein were harmonically restrained during NVT equilibration; during NPT equilibration, only Cα atoms were restrained. The restraint force constant was 1000 kJ/(mol·nm²).

## Data availability

Optimized parameters for carotenoid simulations are provided in Supplementary File 1. Molecular dynamics trajectories were deposited into Zenodo and can be accessed using the following link: https://doi.org/10.5281/zenodo.21716448

## Supplementary Material

Optimized parameters for carotenoid simulations, instructions for using the CARS force field parameters and Supplementary Figures 1-2 are available as Supplementary Materials.

## Acknowledgements

Development of optimized parameters for carotenoid simulations was supported by the Ministry of Science and Higher Education of the Russian Federation (agreement 075-03-2026-305, project FSMG-2025-0003). Molecular dynamics simulations of microbial rhodopsins were supported by the Russian Science Foundation grant 25-74-00077, https://rscf.ru/project/25-74-00077/.

## Author contributions

A.N. initiated the project, developed CARS and prepared the initial draft of the manuscript. Y.O. performed MD simulations of the protein–carotenoid systems. V.K. contributed to validation of lipid parameters. I.G. supervised the project. All authors contributed to preparation of the final version of the manuscript.

