## Supplementary Materials for "CARS: A General Force Field for Carotenoids"

**Supplementary Material:**

**Instructions for Using the CARS Force Field Parameters,**
**Supplementary Figures 1-2**

### Instructions for Using the CARS Force Field Parameters

*Contents of the Parameter Files*

The CARS force field is distributed as two Amber-format frcmod files:

1. `CARS.frcmod` – parameters developed in this work. This file contains the 67 bonded parameters optimized in this study together with the GAFF2 bonded parameters required for carotenoids.
2. `CARS_lipid21.frcmod` – supplementary file containing only the parameters that are taken from Lipid21 for saturated and monounsaturated hydrocarbon chains. This file must be loaded after CARS.frcmod; otherwise the CARS parameters will overwrite the Lipid21 values for alkanes and alkenes.

*Choice of the Force Field Version*

- CARS is recommended for carotenoids that do not contain saturated or nearly saturated lipid tails, including glycosylated carotenoids and molecules with a psi-end group.
- CARS+Lipid21 is recommended for carotenoids carrying saturated or monounsaturated lipid modifications (e.g., fatty acid esters, phytanyl chains). It provides the accuracy of Lipid21 for the lipid moiety while retaining the CARS description of the carotenoid scaffold.

*Charge Assignment and Atom-Type Verification*

All CARS parameters were trained with AM1-BCC charges. Consistency with the training protocol is best achieved by assigning AM1-BCC charges via antechamber (AmberTools23) using the following parameters:

```
antechamber -i carotenoid.mol2 -fi mol2 -o carotenoid_a.mol2
-fo mol2 -c bcc -s 2 -at gaff2
```

Because antechamber also assigns GAFF2 atom types, the following manual checks are required after running the command:

- **Polyene chain:** Verify that single bonds in the polyene chain are assigned as ce–ce or cf–cf, whereas double bonds are assigned as ce–cf. Correct any erroneous assignments by editing the MOL2 file before loading it into tleap.
- **Six-membered sugars:** change the atom type of $sp^3$ carbons in the sugar ring from c6 to c3. This step separates the sugar parameters from the carotenoid ring parameters, as was done in the CARS training set.
- **Triple bonds:** For carotenoids containing C≡C bonds, verify that the **ch** atom type is assigned to the sp carbon adjacent to the ring group and **cg** to the

distal carbon. If necessary, correct the atom types manually in the MOL2 file before loading it into tleap.

Other charge schemes (e.g., RESP, EspalomaCharge) have not been validated with CARS and should be used with caution.

*Building a System with tleap*

Before assembling the system, run parmchk2 to generate an `.frcmod` file containing any additional bonded parameters that are required for the molecule but are not present in the CARS parameter set:

```
parmchk2 -i carotenoid_a.mol2 -f mol2 -o missing.frcmod
```

A minimal working example for a single carotenoid is given below.

```
source leaprc.gaff2

loadamberparams missing.frcmod
loadamberparams CARS.frcmod
loadamberparams CARS_lipid21.frcmod   # only for CARS+Lipid21

car = loadmol2 carotenoid_a.mol2

saveamberparm car system.prmtop system.inpcrd
quit
```

This procedure yields a fully parameterised system that can be converted to the GROMACS format with ParmEd or used directly in Amber.

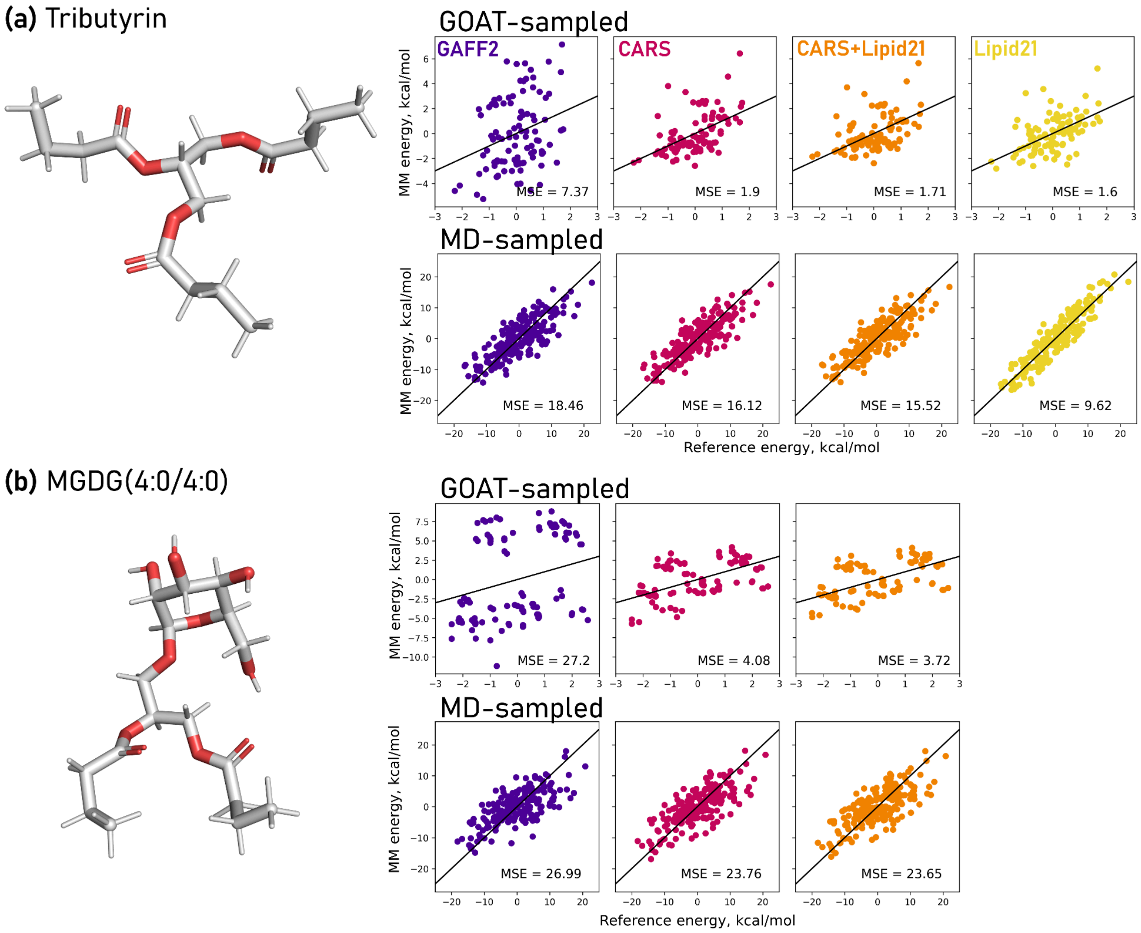


**Figure S1.** Comparison of reference QM (r2SCAN-3c) and force-field energies for tributyrin (a) and MGDG(4:0/4:0) (b). For each molecule, the structure is shown alongside the scatter plots for MD-sampled (upper row) and GOAT-sampled (lower row) conformers. Energies were computed with GAFF2, CARS, CARS+Lipid21, and, for tributyrin only, Lipid21. The black diagonals represent perfect agreement between the energy values.

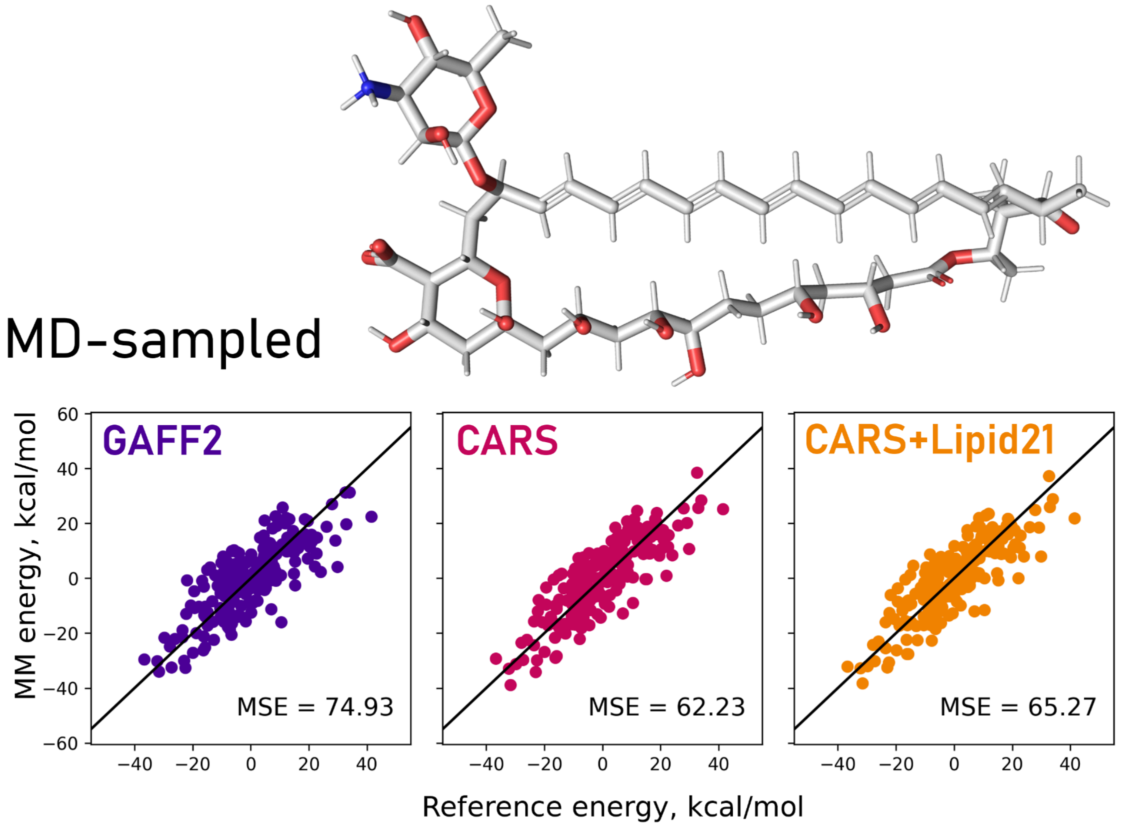


**Figure S2.** Comparison of reference QM (r2SCAN-3c) and force-field energies for amphotericin B. The structure is shown alongside the scatter plots for MD-sampled conformers. Energies were computed with GAFF2, CARS and CARS+Lipid21. The black diagonals represent perfect agreement between the energy values.
